# Pair-EGRET: enhancing the prediction of protein-protein interaction sites through graph attention networks and protein language models

**DOI:** 10.1101/2023.12.25.572648

**Authors:** Ramisa Alam, Sazan Mahbub, Md. Shamsuzzoha Bayzid

**Affiliations:** Department of Computer Science and Engineering Bangladesh University of Engineering and Technology Dhaka-1205, Bangladesh; Computational Biology Department, School of Computer Science Carnegie Mellon University, Pittsburgh, PA 15213, USA

**Keywords:** protein-protein interaction sites, deep learning, graph neural network, attention mechanism, protein language model

## Abstract

Proteins are responsible for most biological functions, many of which require the interaction of more than one protein molecule. However, accurately predicting protein-protein interaction (PPI) sites (the interfacial residues of a protein that interact with other protein molecules) remains a challenge. The growing demand and cost associated with the reliable identification of PPI sites using conventional experimental methods call for computational tools for automated prediction and understanding of PPIs. Here, we present Pair-EGRET, an edge-aggregated graph attention network that leverages the features extracted from pre-trained transformer-like models to accurately predict PPI sites. Pair-EGRET works on a *k*-nearest neighbor graph, representing the three-dimensional structure of a protein, and utilizes the cross-attention mechanism for accurate identification of interfacial residues of a pair of proteins. Through an extensive evaluation study using a diverse array of experimental data, evaluation metrics, and case studies on representative protein sequences, we find that our method outperforms other state-of-the-art methods for predicting PPI sites. Moreover, Pair-EGRET can provide interpretable insights from the learned cross-attention matrix. Pair-EGRET is freely available in open source form at (https://github.com/1705004/Pair-EGRET).

## 1 Introduction

Proteins play a fundamental role in various cellular processes necessary for the survival of living organisms. Many of these processes rely on the formation of protein complexes, which involve interactions between two or more proteins [1]. A thorough understanding of protein interfaces and the interacting residues involved is crucial in fields like disease research, drug development, and unraveling the underlying mechanisms of basic cellular biology [2–7].

Protein-protein interaction sites (PPIS) are the residues of a protein that interact with other proteins’ residues to form an interface between them. There are primarily two formulations for the problem of identifying interacting residues in protein complexes. One is *partner-independent* prediction problem that tries to find the set of residues of a given isolated protein that may interact with residues of any other protein [8, 9]. The second form – and the one addressed in this study – is *partner-specific* which involves the identification of interacting residues of a protein for a particular partner protein. Partner-specific methods can further be classified into those that seek to only identify which residues are part of the interface between the protein pairs (henceforth referred to as interface region prediction methods) and those that seek to predict which specific pairs of residues (one from each protein) interact with one another (henceforth referred to as pairwise PPIS prediction methods) [10, 11]. The latter is more challenging than the first and for most methods, performing the pairwise PPIS prediction task leads to the identification of interface regions. This research addresses both forms, presenting an innovative method for accurately predicting interface regions and pairwise PPIS in protein complexes.

Experimentally identifying PPIS using wet lab methods is time-consuming and costly, resulting in the rise of various computational approaches as alternatives. Computational methods like protein-protein docking models [12–15], template-based methods [16–18] and machine-learning methods [11, 19–24] employ different techniques like computationally rotating and translating proteins to produce different poses, comparing an unknown protein with a known query protein, and training models to learn features of interacting residues, etc. However, these methods suffer from numerous shortcomings including limited coverage, inability to predict novel interactions, and challenges in feature selection.

Some recent deep learning-based methods [8, 10, 25, 26] have demonstrated that incorporating information from both the primary amino-acid sequence and the 3D structure of a protein leads to more accurate identification of PPIS. For extracting features from the primary sequence, the recently developed protein language models [27, 28], trained on large datasets of 1D amino acid sequences, have been proved to be effective [8,26,29]. For encoding the 3D structural information of proteins, several geometric structures have been proposed [9,30] among which Graph Neural Networks (GNN) [31] has proved to be useful for a number of methods [8, 10, 25]. GNNs have the capability to learn global and local contextual features for a residue from its neighborhood. Mahbub and Bayzid [8] proposed an **E**dge Aggregated **GR**aph Attention N**ET**work (EGRET), a variant of GNN [32], where both node-level and edge-level features are used for calculating the attention scores allowing the model to utilize the rich structural data encoded in the edges of the graph. EGRET was proposed for the task of partner-independent prediction of interaction sites.

In this study, building on the recent successful application of transformers and GAT networks in partner-independent PPI site prediction, we propose a novel deep learning model Pair-EGRET that extends the architecture of EGRET for both pairwise PPI site and interface region predictions. Pair-EGRET integrates GNN and protein language models to effectively leverage both structural and sequence-based information. Moreover, to allow each protein to attend to the relevant features from the other protein’s residues, we adopted the concept of the cross-attention [33], widely used in Natural Language Processing (NLP) for incorporating information from multiple input sources or contexts. The combination of using transfer learning, edge-aggregated GAT networks, and cross-attention modules led to the improvement of Pair-EGRET compared to the best alternative methods on widely accepted benchmark datasets for partner-specific interaction prediction (both PPI site and interface region predictions). We have included case studies that visually inspect the predicted binding residues. We further visualized and interpreted the representations learned by Pair-EGRET. Pair-EGRET is a fully automated end-to-end pipeline, alleviating the need for time-consuming and careful hand-tuning.

## 2 Approach

### 2.1 Feature Representation

We represent the three-dimensional structures of the receptor and the ligand of a complex using directed *k*-nearest neighbor graphs [34], denoted as *G*_*receptor*_ and *G*_*ligand*_ respectively. Each graph node corresponds to an amino acid residue and is connected to its *k* closest neighbors via directed edges, where *k* is a hyperparameter. The determination of nearest neighbors is based on averaging the distances between the atoms of residue pairs within a protein, which are obtained from Protein Data Bank (PDB) files [35]. We used embedding vectors generated by ProtBERT [27] and some physicochemical properties of amino acids as node-level features. The distance and angle between residues served as the edge-level features for the weighted edges connecting them.

#### Node-level features

Each residue *i* in a protein is associated with a feature vector 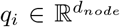, where *d*_*node*_ represents the number of node features used in this study. We used two types of node features to represent a residue.

1. **Embedding-based features** of the residues were extracted from the protein sequences using Prot-BERT, a contextual embedding generation pipeline developed by [27]. ProtBERT captures both local and global context, including neighboring residues and overall protein structure, to generate embedding vectors *e* = *e*_1_, *e*_2_, …, *e*_*N*_, 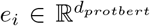 (*d*_*protbert*_ = 1024 and *N* = number of residues in the given protein), which encode the structural and functional characteristics of the residues. Note that other embedding generation models available in ProtTrans [28], such as ProtXL or ProtXLNet, can also be used instead of ProtBERT.
2. **Physicochemical Features** of amino acids were incorporated as node features. These features encompass a range of properties, including hydrophilicity, flexibility, accessibility, turns scale, exposed surface, polarity, antigenic propensity, hydrophobicity, net charge index of side chains, polarizability, solvent-accessible surface area (SASA), relative SASA, side-chain volume, and residue depth. Notably, we calculate relative hydrophobicity and polarity based on two different scales or methods, namely H11a, H12a, P11a, and P12a, respectively, to ensure a comprehensive representation of these characteristics in our analysis [36]. These properties of a node *i* is represented by a vector 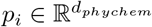 (*d*_*phychem*_ = 16). By concatenating the vectors *e*_*i*_ and *p*_*i*_ we obtained the final node features *q* = {*q*_1_, *q*_2_, …, *q*_*N*_}, 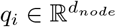 where *d*_*node*_ = *d*_*protbert*_ + *d*_*phychem*_ = 1024 + 16 = 1040.

#### Edge-level features

Similar to EGRET [8], the edge features of an edge from node *j* to node *i* in the graph representation of a protein is denoted by *ξ*_*ji*_, where 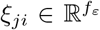 and *f*_*ε*_ is the number of features of the edge. We used the following two features (i.e., *f*_*ε*_ = 2) as edge-features: (i) inter-residue distance *D*_*ij*_ which denotes the average distance between the atoms of the residues, and (ii) relative orientation *θ*_*ij*_ which is measured by the absolute value of the angle formed by the surface-normals of the planes passing through the alpha carbon atom (*C*_*α*_), the carbon atom of the Carboxyl group, and the nitrogen atom of the Amino group of each residue.

### 2.2 Architecture of Pair-EGRET

The architecture of Pair-EGRET can be discussed in two parts: (i) the architecture of EGRET [8], which was proposed for predicting the interaction sites of a single isolated protein and is used as the foundation of Pair-EGRET, and (ii) the proposed extension of the EGRET architecture for predicting interaction sites from pairs of proteins.

#### 2.2.1 Architecture of EGRET

We briefly discuss the architecture of EGRET here to make the paper self-contained and easy to understand. The core components of the EGRET model are (i) the local feature extractor, (ii) the edge-aggregated graph attention layer, and (iii) the node-level classifier.

##### Local feature extractor

The local feature extractor captures local interactions of the protein residues with other “sequentially closer” residues (not necessarily close in Euclidean space) while reducing the dimensionality of the node-level features. A one-dimensional convolutional neural network with a small odd number window size is used to encode the node feature vectors *q* = {*q*_1_, *q*_2_, …, *q*_*N*_} into a new condensed and neighbor-aware feature representation *h* = {*h*_1_, *h*_2_, …, *h*_*N*_}, 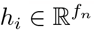, where *f*_*n*_ *< d*_*protbert*_.

##### Edge-aggregated graph attention layer

The edge-aggregated graph attention layer transforms the features *h*_*i*_ of the node *i* by encoding the three-dimensional structural information of its neighborhood *N*_*i*_. This layer uses a modified version of the original graph attention layer [32] and aggregation process used in various GNN-based architectures [32, 37]. In the original aggregation process, the node features are transformed by taking a weighted average of the neighborhood node features using the equation: 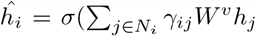) where, 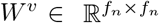 is a learnable parameter and *γ*_*ij*_ is the attention score calculated from *h*_*i*_ and *h*_*j*_ that represents the importance of the features of node *j* to node *i*. EGRET improves upon this method by incorporating edge features during the calculation of attention scores and the aggregation process, resulting in a new scoring function *e*_*ji*_ and attention distribution *α*_*ji*_. These metrics are obtained from the following equations.

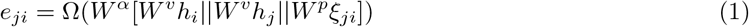

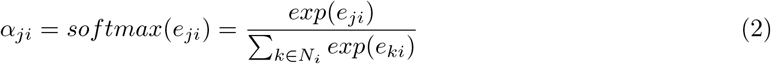

Finally, the node and edge features are aggregated using the equation

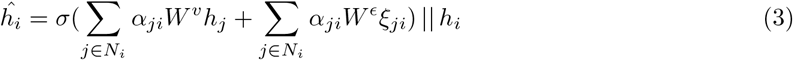

Here, 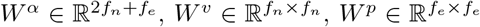 and 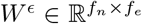 are learnable parameters, ∥ is the concatenation operator, and Ω (.) and *σ* (.) are activation functions.

##### Node level classifier

This final layer linearly transforms the aggregated features obtained from the previous layer *ĥ*_*i*_ and applies sigmoid activation to generate interaction probabilities for each residue of a sequence.

Figure 4 in Appendix A shows the overall end-to-end pipeline of EGRET. It demonstrates EGRET being applied to a dummy protein with 13 residues.

#### 2.2.2 Extension of EGRET for pairwise prediction

In this study, we extend the EGRET architecture to identify interactions between protein pairs. Since the original EGRET was developed for single isolated proteins, we leverage its architecture for extracting useful features from the graph representations of the receptor and the ligand separately, which are subsequently analyzed using a multi-headed cross-attention layer.

The core components of the pair-EGRET architecture include (i) Siamese EGRET network, (ii) positional encoder, (iii) multi-headed cross-attention layer, (iv) pairwise classifier, and (v) interface region classifier. The first three components are connected sequentially and are common to the architecture required for both the problems we are addressing in this study, i.e. pairwise interaction site and interface region prediction. The final two layers (pairwise classifier and interface region classifier) are parallel networks that generate the outputs corresponding to these two problems. The overall end-to-end pipeline of Pair-EGRET is shown in Figure 1, which demonstrates the major modules of Pair-EGRET being applied to a pair of proteins each containing 13 residues.

**Figure 1:**
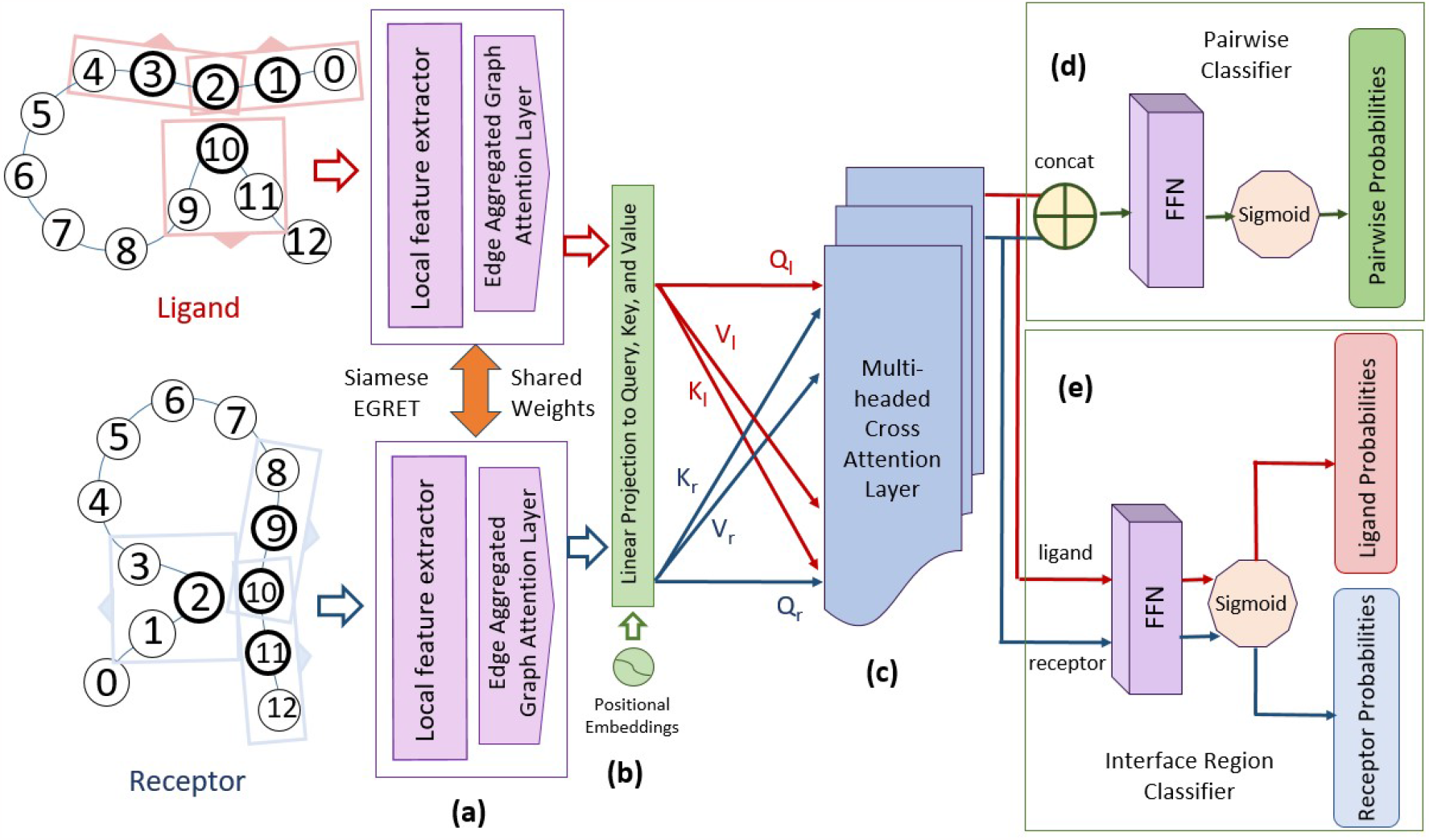
Schematic diagram of the overall pipeline of Pair-EGRET being applied to a receptor protein (r) and a ligand-protein (l). **(a)** Siamese EGRET network with shared weights. **(b)** Positional encoder being added to the individual proteins and linearly projected to three separate feature spaces– query, key, and value. **(c)** Both proteins transformed through the multi-headed cross-attention layer using the query vector of itself and the key and value vector of the other protein. **(d)** Pairwise classifier applied to the merged features of both proteins to produce pairwise probabilities of interaction. **(e)** Interface region classifier applied to individual proteins for producing probabilities of being part of the interface for each residue.

##### Siamese EGRET network

The architecture of Pair-EGRET begins with a Siamese network, which consists of a pair of identical networks containing the first two modules of EGRET (local feature extractor and edge-aggregated graph attention layer). These two networks work in tandem on the given pair of proteins (the receptor and ligand) and share their weights, enabling the Siamese network to learn a common representation of both the ligand and receptor, ensuring that the model learns a common representation of proteins and is invariant to the ordering of the partner proteins provided to it.

Given that the third module of EGRET (the node-level classifier) calculates the interaction probability for each residue independently of its partner protein in the complex, we eliminate this layer. Instead, we utilize features extracted from EGRET’s local feature extractor and GAT layer to encode each residue with features derived from its neighborhood, taking into account both sequential and spatial proximity.

The node features *q*_*i*_ and edge features *ξ*_*ij*_ corresponding to each of the graphs *G*_*receptor*_ and *G*_*ligand*_ are fed through the Siamese network to generate features *ĥ*_*l*_ and *ĥ*_*r*_ respectively. In addition to the embedding-based features used as node features in the original EGRET framework, we integrate a wide array of physicochemical features specific to each residue. Furthermore, we enhance the quality of the features generated from the Siamese network by increasing the number of convolution layers in the local feature extractor and using multiple layers of the graph attention module.

##### Positional encoder

The positional encoder module – inspired by [33] – is used in Pair-EGRET to utilize the information about the position of residues in a protein sequence. This module calculates a positional embedding for each residue and adds it to the feature representations obtained from the Siamese network. This can be useful because the location of a residue can impact its interaction with residues present in the partner protein in the complex. Following Vaswani et al., [33] we defined the positional encoding for a position *pos* in the sequence as follows:

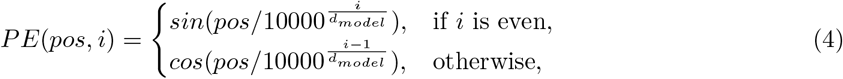

where *i* ∈ [1, *d*_*model*_] is the dimension of the embedding vector being calculated, and *d*_*model*_ is the dimension of the input embedding vector. This function maps the position of each amino acid to a set of position-specific embeddings *PE. PE* is added to the feature representations *ĥ*_*l*_ and *ĥ*_*r*_ obtained from the siamese network, to generate the outputs *P* ^*l*^ and *P* ^*r*^ of this layer.

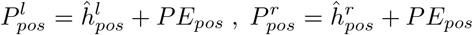

##### Multi-headed cross-attention layer

The multi-headed cross-attention layer employs the cross-attention mechanism [33] to combine features originating from both receptor and ligand proteins. Through an encoder-decoder structure, this layer facilitates the mutual exchange of information between the proteins. It quantifies the influence of a ligand residue on a receptor residue and vice versa by generating attention scores that help identify relevant residue pairs within a complex. We employ multiple attention heads within this module to capture various aspects of the input features.

The cross-attention module transforms the input feature vectors into three feature spaces: query, key, and value using learnable parameters *W*_*Q*_, *W*_*K*_, and 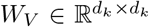, where *d*_*k*_ is a hyperparameter. Specifically, the ligand feature vector *P* ^*l*^ obtained from the positional encoder layer is transformed into query 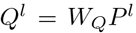, key 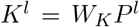, and value 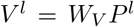, while the receptor feature vector *P* ^*r*^ is transformed into *Q*^*r*^, *K*^*r*^, and *V* ^*r*^ using similar equations.

To model the interaction between the proteins, the cross-attention module is applied to the ligand and receptor feature vectors separately, as defined by the following equations:

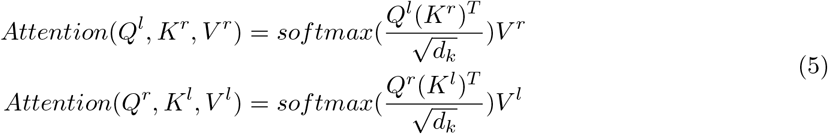

where *softmax* is the softmax activation function, and (*·*)^*T*^ denotes the transpose operation. To capture diverse aspects of the input features, the cross-attention module employs multiple attention heads. Each attention head is an independent attention layer that attends to different aspects of the input. Specifically, multi-headed attention is defined by the following equation:

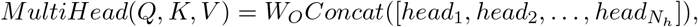

where *head*_*i*_ = *Attention*(*Q, K, V*) is the output of the *i*-th attention head, *N*_*h*_ is the number of total attention heads, and *W*_*O*_ is a learnable parameter used to project the concatenated attention heads back to the original feature space.

The outputs of the multi-headed attention are then added to the input of the module through a residual connection [38] which enables the model to capture both the attended features and original features, followed by layer normalization [39] which stabilizes the model. The final outputs *M* ^*l*^ and *M* ^*r*^ are defined by the following equations:

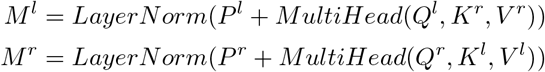

##### Pairwise classifier

The pairwise classifier layer in Pair-EGRET produces the final output for our first task: pairwise PPIS prediction. For each residue pair (*l*_*i*_, *r*_*i*_), this module generates an output *O*_*i*_ *∈ {*0, 1*}* where *O*_*i*_ = 1 would indicate that the residue *l*_*i*_ in the ligand and the residue *r*_*i*_ in the receptor are an interacting pair, whereas *O*_*i*_ = 0 would indicate otherwise.

For each residue pair, we extract the node features 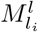 and 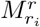 from the outputs *M* ^*l*^ and *M* ^*r*^ generated by the cross-attention module and concatenate them. This representation of the residue pair is passed through a feed-forward network (FFN) that incorporates a combination of fully connected layers, nonlinear activation functions such as ReLU and Leaky ReLU, and finally, a Sigmoid activation function to predict the probability of interaction between the nodes *l*_*i*_ and *r*_*i*_.

Similar to [10], to ensure invariance to the ordering of the receptor and ligand, we concatenate the features in both possible orders, resulting in two predictions:

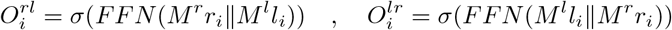

where *FFN* is the feed forward network and *σ* is the Sigmoid activation function. Finally, we take the average of these two predictions to generate the final output probability *O*_*i*_ for the residue pair (*l*_*i*_, *r*_*i*_):

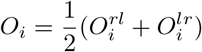

##### Interface region classifier

The interface region classifier generates the final outputs for our second task: identifying the interface region of protein complexes. For each residue *l*_*i*_ of the ligand or *r*_*j*_ of the receptor this module generates an output 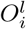 or 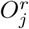 which indicates whether the residues is a part of the interface of the corresponding complex or not.

In this layer, instead of concatenating the outputs *M* ^*l*^ and *M* ^*r*^ generated by the cross-attention layer, we pass *M* ^*l*^ and *M* ^*r*^ individually through a feed-forward network (FFN). Similar to the pairwise classifier, this network incorporates fully connected layers and activation functions that linearly transform the feature vectors and finally a Sigmoid activation function generates the probabilities of each residue being part of the interface region:

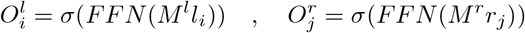

## 3 Experimental Studies

We conducted a comprehensive evaluation study to compare the performance of Pair-EGRET, a novel method proposed in this study, with state-of-the-art conventional machine learning methods, convolutional neural network (CNN)-based methods, and graph neural network (GNN)-based methods, using widely-accepted benchmark datasets for pairwise interaction site and interface region prediction.

### 3.1 Dataset

We evaluated Pair-EGRET on three benchmark datasets: i) Docking Benchmark version 5.0 (DBD5) [40], ii) Dockground X-ray unbound benchmark [41], and iii) A subset of the MaSIF [9] dataset. For each complex, these datasets contain the atomic coordinates of the amino acid residues of the proteins in both their bound and unbound forms. We used the unbound structures to produce the input features for the model and the bound structures to identify interacting residues in protein pairs. Consistent with prior work [10, 11, 42], we defined two residues to be interacting if any of their non-Hydrogen atoms were within 6Å of one another.

We used the DBD5 and Dockground datasets for evaluating Pair-EGRET and other contemporary methods on the task of pairwise PPIS prediction. DBD5 was also used along with the MaSIF dataset for the interface region prediction task. The choice of datasets in each case was made based on the availability of evaluation scores of existing methods on the intended task so that a fair and comprehensive analysis of performance could be presented.

i. **Docking Benchmark version 5.0 (DBD5)** is widely recognized as the standard benchmark for evaluating pairwise PPIS prediction and interface region identification. The dataset includes structures of 230 complexes from the protein data bank (PDB) [35] with amino acid sequence lengths of the constituent proteins varying from 29 to 2128. Training and validation on DBD5 were performed using the 175 complexes present in version 4.0 of Docking Benchmark (DBD4) [43]. We performed an 80%-20% partition of the 175 complexes stratifying them by the difficulty provided in [40]. For testing, we used a set of 55 complexes that were added in the update from DBD4 to DBD5. This time-based split of the dataset simulates the ability of the model to predict unreleased complexes, as opposed to a random split which has more training/testing cross-contamination. [44]
ii. **Dockground** is another benchmark used for evaluating pairwise interaction site prediction models. The dataset contains a diverse array of protein complexes of varying difficulties. Compared to DBD5, it has fewer proteins with rigid bodies and more with higher difficulty levels. In our experiments, we used the unbound docking benchmark set 4 of the Dockground dataset containing 396 complexes with only 77 complexes shared with DBD5. We used 236 complexes for training, 60 complexes for validation, and 100 complexes for testing Pair-EGRET on the task of pairwise interaction site prediction.
iii. **(iii) MaSIF** is a relatively large dataset containing a total of 3362 complexes taken from the PRISM [45] list of nonredundant proteins, the ZDock benchmark [13], PDBBind [46], and SabDab [47] dataset. We used the curated subset of MaSIF used by PInet [30] which excludes complexes with interface regions smaller than 1% of the size of the ligand. We also excluded complexes with receptor or ligand sequence lengths smaller than the minimum neighborhood size required by the edge-aggregated graph attention layer of EGRET. This resulted in a dataset containing 3147 complexes, which was split into 1890 training, 470 validation and, 787 test complexes. We used this subset of MaSIF for evaluating Pair-EGRET in identifying interface regions of complexes. This benchmark uses bound conformations of proteins to produce features for training the models. Consistent with other methods, only for this benchmark, we used the provided bound conformations for generating the node and edge-level features of Pair-EGRET. In all of these datasets, the number of positive samples (interacting pairs) is very small compared to the number of negative samples (non-interacting residue pairs) with the ratio of positive-to-negative samples being close to almost 1:1000. To address this imbalance, similar to [10, 25], during training, we downsampled the negative samples to obtain a 1:10 ratio of positive-to-negative samples. For validation and testing, we preserved the original ratio of the samples present in the datasets. The summary of the dataset sizes and data partitions are presented in Appendix B (Table 3).

### 3.2 Performance Evaluation

As the datasets we are using are highly imbalanced, accuracy is not a meaningful evaluation metric for either of the tasks. We used AUROC (Area under receiver operating characteristic curve) and AUPRC (Area under precision-recall curve) as our primary evaluation metrics. AUROC and AUPRC are threshold independent and therefore, are appropriate measures of performance for evaluating performance on high-imbalance datasets [48]. We calculated the AUROC for each of the test complexes and compared the performance of the methods using the median AUROC scores. This reduces the probability of extreme change in the performance resulting from very small or very large complexes. [25]. We also compared the performance of Pair-EGRET in terms of AUPRC, even though the extreme class imbalance resulted in poor AUPRC scores for all the methods in the pairwise interaction site prediction task.

#### 3.2.1 Pairwise PPIS prediction results

##### Results on DBD5

For the task of pairwise interaction site prediction, we compared the median AUROC and AUPRC scores of Pair-EGRET with the state-of-the-art machine learning, CNN, and GNN-based methods [10, 11, 44, 49–53]. It is evident from Table 1 that Pair-EGRET outperforms all the other methods in terms of our primary evaluation metric, median AUROC with a score of 0.88828. We note that all the methods achieved low AUPRC scores, indicating the complexity of the task and the imbalanced nature of the dataset. However, our method achieves an AUPRC score of 0.0173 which is comparable to the best-performing method DCNN [49] with an AUPRC of 0.018.

**Table 1:** Comparison between the predictive performance of different methods in predicting pairwise PPIS of the test complexes of DBD5 and Dockground. Scores for the baseline methods on DBD5 are reported from [10, 25, 26]. Values not reported by corresponding studies are indicated by ‘-’.

| DBD5 |  |  |  |  |  | Dockground |  |
| --- | --- | --- | --- | --- | --- | --- | --- |
| Method | Median AUROC | AUPRC | Method | Median AUROC | AUPRC | Method | Median AUROC |
| BIPSPI | 0.878 | - | DTNN | 0.867 | 0.007 | BIPSPI+ | 0.831 |
| SASNet | 0.876 | - | NEA | 0.876 | 0.012 | Pair-EGRET | **0.8747** |
| DCNN | 0.828 | **0.018** | EGNN | 0.829 | - |  |  |
| NGF | 0.865 | 0.007 | GVP-GNN | 0.885 | - |  |  |
| Pair-EGRET | **0.888** | 0.0173 |  |  |  |  |  |

##### Results on Dockground

Although the unbound benchmark version 4 of Dockground is less explored for the pairwise PPIS prediction task, we evaluated Pair-EGRET on this benchmark because of the varying levels of difficulty it offers making it more diverse than DBD5. Compared to BIPSPI+ [54] – one of the few methods evaluated on Dockground for this task – Pair-EGRET performs substantially better with a median AUROC of 0.8747, highlighting its robustness in identifying interaction sites in relatively difficult complexes.

#### 3.2.2 Interface region prediction results

##### Results on DBD5

We compared Pair-EGRET with BIPSI, BIPSPI+, and PInet [30] for interface region prediction on the DBD5 test dataset. In addition to AUROC and AUPRC, we also report the precision and recall scores of the methods for a fair comparison and consistency with the other prior studies that analyzed this dataset. Pair-EGRET substantially outperforms all the other methods under two evaluation metrics-AUROC and recall. Remarkably, the AUROC score of Pair-EGRET is 0.924 which is 8.96% higher than the second best method BIPSPI+. However, Pair-EGRET performed worse than other methods in terms of precisions and AUPRC on this particular dataset.

**Table 2:** Performance evaluation of different methods in identifying interface regions of test complexes of DBD5 and MaSIF datasets. geom* indicates models that only use geometric features of proteins. Values that were not reported in the MaSIF study [30] are indicated by ‘-’.

| DBD5 |  |  |  |  | MaSIF |  |  |
| --- | --- | --- | --- | --- | --- | --- | --- |
| Method | AUROC | AUPRC | Precision | Recall | Method | AUROC | AUPRC |
| BIPSPI | 0.822 | 0.410 | 0.391 | 0.558 | SPPDER | 0.65 | - |
|  |  |  |  |  | MaSIF geom* | 0.68 | - |
| BIPSPI+ | 0.848 | 0.4653 | 0.438 | 0.573 | MaSIF | 0.87 | - |
|  |  |  |  |  | PInet geom* | 0.75 | 0.30 |
| PInet | 0.753 | **0.596** | **0.492** | 0.723 | PInet | 0.88 | 0.45 |
| Pair-EGRET | **0.924** | 0.275 | 0.255 | **0.746** | Pair-EGRET | **0.9583** | **0.5938** |

##### Results on MaSIF

Our assessment of Pair-EGRET on the MaSIF dataset involved a comparison with MaSIF [9], SPPIDER [55], and PInet [30]. Pair-EGRET significantly outperformed all other methods in terms of both AUROC and AUPRC. Pair-EGRET achieved an AUROC score of 0.9583, which is 8.89% than the next best method PInet. Additionally, Pair-EGRET achieved a much higher AUPRC score (0.5938) which is around 15% higher than PInet. Since the MaSIF benchmark is substantially large and contains a collection of diverse and uncurated protein complexes [30], the substantially higher AUROC and AUPRC scores of Pair-EGRET than other methods are indicative of the model’s ability to learn a better-generalized representation of interface residues of ligand and receptor proteins compared to other methods.

### 3.3 Interpretability analysis: patterns of cross-attention scores

One of the noteworthy contributions of this study is introducing the cross-attention mechanism for pairwise protein interaction site and interface region prediction. We investigated the behavior of the multi-headed cross-attention layer of Pair-EGRET to provide insights into how this layer learns the correlation between residues from two different proteins. Figure 2(c) shows the heatmap representation of the attention scores generated for a representative protein (PDB ID 3HI6) from the DBD5 test dataset. For the convenience of visualization, we show a small window of the attention score matrix corresponding to 20 residues (residue IDs 128-147) from chain A of the ligand and 20 residues (residue IDs 22-41) from chain L of the receptor.

**Figure 2:**
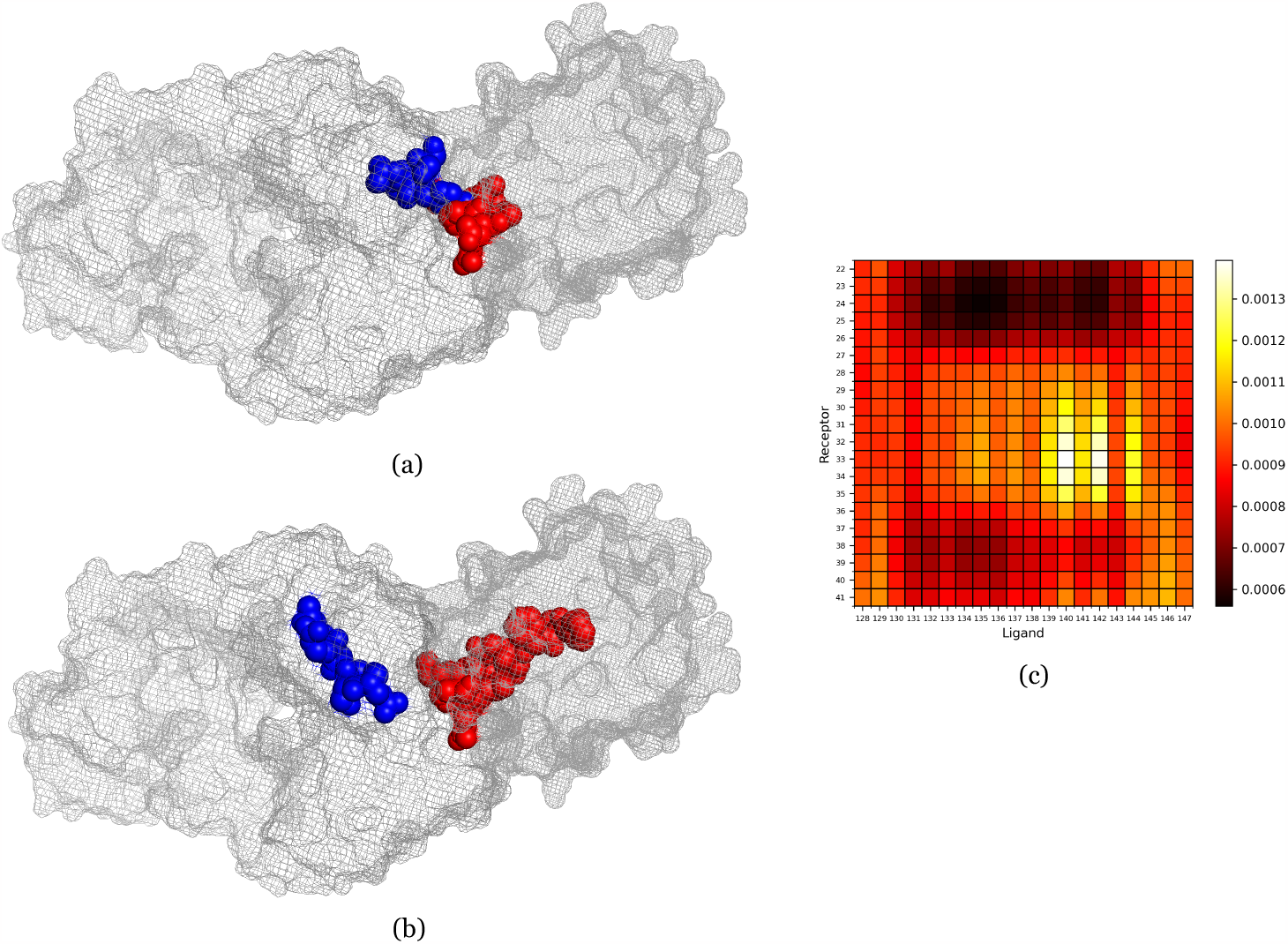
Patterns of cross-attention scores in a 20 residue window of chain A of the ligand and chain L of the receptor of a representative complex (PDB ID 3HI6). **(a)** PyMOL visualization of the residues corresponding to the lighter regions (high attention scores) of the heatmap. **(b)** PyMOL visualization of the residues corresponding to the darker regions (low attention scores) of the heatmap. **(c)** Heatmap of the attention scores generated by the multi-headed cross-attention layer of Pair-EGRET.

Interestingly, the residue pairs that were given the highest attention scores (lighter color in the heatmap) either correspond to interacting residue pairs (according to the dataset labels) or are very close to such interacting pairs in the Euclidean space. As shown in the PyMOL [56] visualization of Figure 2(a), the residues having relatively high attention scores are very close in space and likely belong to the interface region of the protein complex. On the other hand, pairs with lower attention scores (darker color in the heatmap) correspond to the residues shown in Figure 2(b). These residue pairs are visibly apart from each other and some are even inside the surface of the proteins. These results demonstrate the meaningful relationship between attention scores generated by the cross-attention layer of Pair-EGRET and the characteristics of interacting residue pairs in a complex.

## 4 Case study

In Figure 3, we visualized two representative complexes (PDB ID 3HI6 and 1JTD) from the DBD5 test dataset using the PyMOL software in order to perform a qualitative analysis of Pair-EGRET’s performance in identifying interface regions. We also visualized the interface regions predicted by NEA (pairwise PPISP prediction method developed by Fout et al. [10]) to compare our results with a competitive method. Most of the existing methods are not publicly available (either as user-friendly software packages or web servers). Consequently, we were unable to include additional methods in this visual inspection.

**Figure 3:**
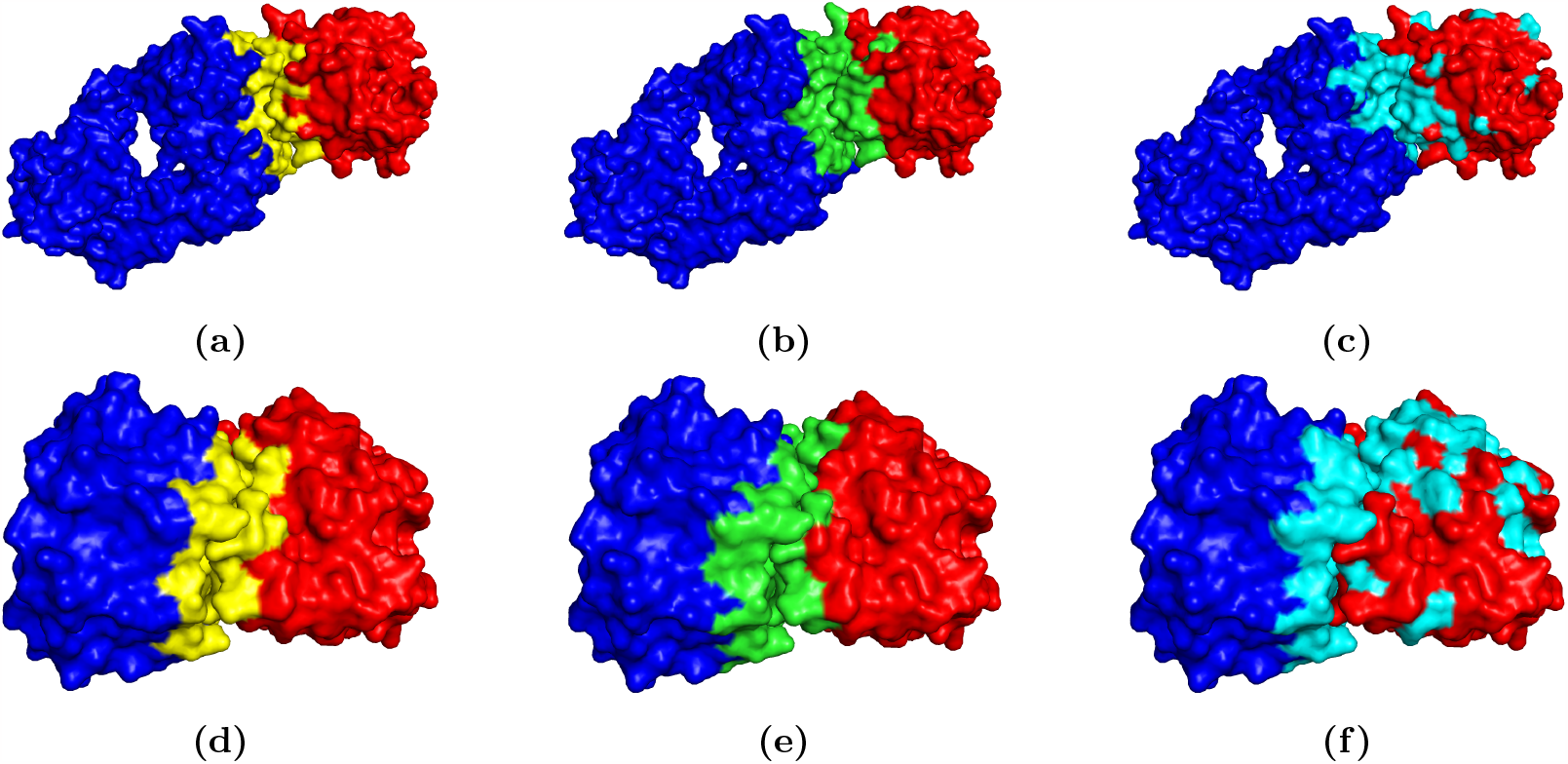
PyMOL visualization of two representative complexes (PDB ID 3HI6 and 1JTD) from the test set of DBD5 in bound form. The red and blue surfaces represent the ligand and receptor proteins respectively. **(a-c)** The true (yellow), predicted by Pair-EGRET (green), and predicted by NEA (cyan) interface regions of 3HI6. **(d-f)** The true (yellow), predicted by Pair-EGRET (green), and predicted by NEA (cyan) interface regions of 1JTD.

It is evident that compared to the predictions generated by NEA (Figure 3c, 3f), the interface residues identified by Pair-EGRET (Figure 3b, 3e) are less scattered and more concentrated towards the true interface regions (Figure 3a, 3d). Moreover, predictions from Pair-EGRET accurately encompass the entirety of the actual interface whereas NEA misses parts of the interface regions in both complexes. Furthermore, Pair-EGRET produced notably lower numbers of false negative predictions than NEA.

## 5 Conclusions

We presented Pair-EGRET, a novel deep-learning method for accurately identifying pairwise interaction sites and interface regions of protein complexes. We have demonstrated the effectiveness of using an edge-aggregated graph attention network as well as cross-attention mechanism in the context of pairwise interaction prediction. Our systematic analyses of the performance of different methods under various model conditions indicate the predictive power and effectiveness of Pair-EGRET in pairwise PPI site and interface region predictions. Furthermore, Pair-EGRET offers a more interpretable framework than the typical black-box deep neural network methods.

This study can be extended in several directions. We utilized the protein language model Prot-Trans [28] for feature generation. A more recent and larger language model (ESM-2) [57] for proteins, released by Facebook Research, can be utilized for producing better predictions by Pair-EGRET. The increasing availability of structure-known proteins has led to the assembly of several larger datasets [9,44] that can be used for further traininge our model leading to enhanced performance.

## Appendix A Architecture of EGRET

**Figure 4:**
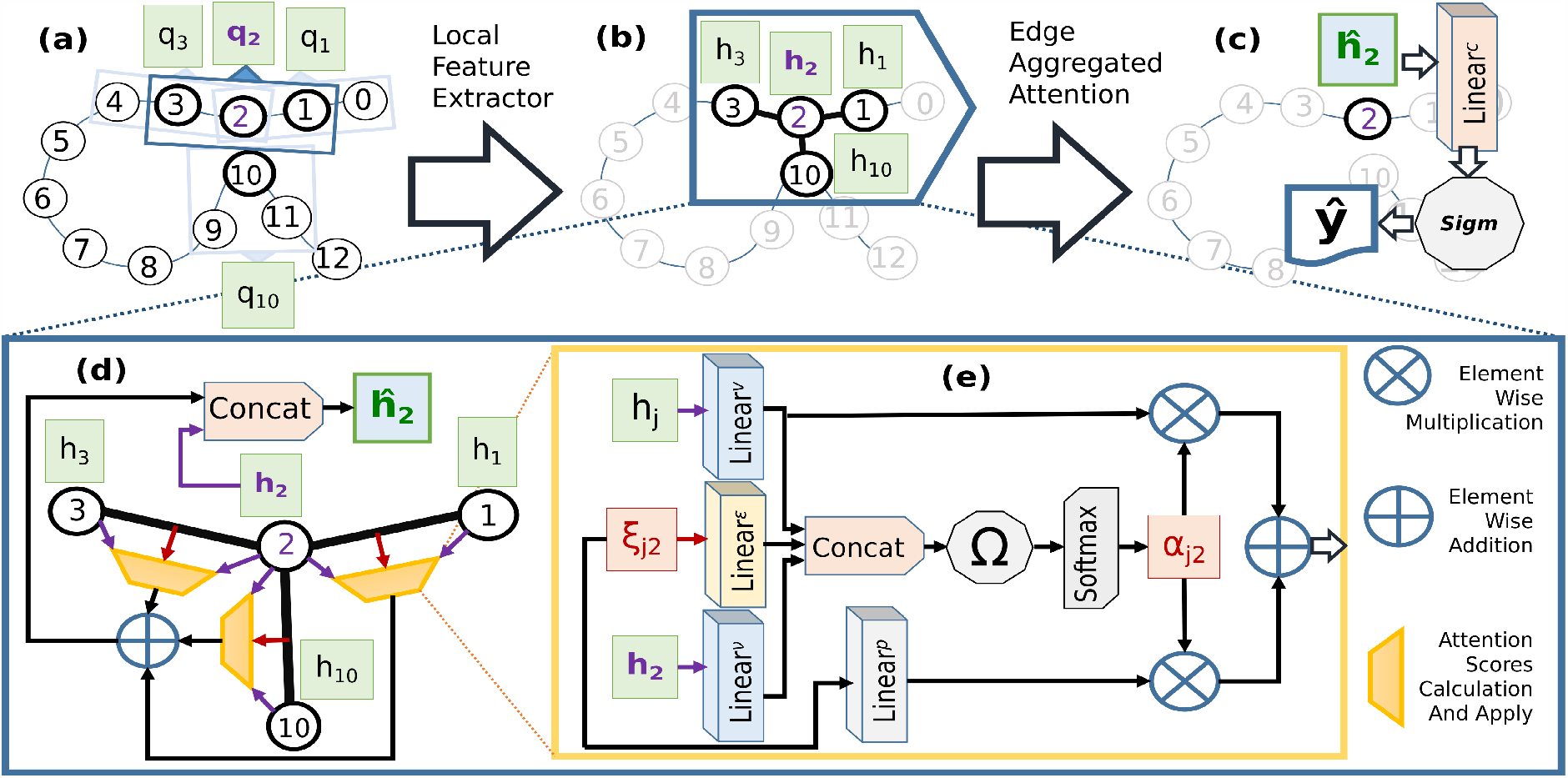
Schematic diagram of the overall pipeline of EGRET being applied to a dummy protein having 13 residues. **(a)** Local feature extractor (with window size *w*_*l*_*ocal* = 3). **(b)** Edge-aggregated graph attention layer applied to residue 2 with neighborhood *N*_2_ = *{*1, 3, 10*}*. **(c)** Node level classifier applied to final representation *ĥ*_2_ of node 2. **(d)** The details of the edge-aggregated graph attention layer in an expanded form. **(e)** The expanded form of the module that calculates the attention scores for aggregation. This figure has been taken from [8].

## Appendix B Summary of Dataset

**Table 3:** Summary of the datasets used in this study.

| Dataset | Samples | Train | Validation | Test | Total |
| --- | --- | --- | --- | --- | --- |
| DBD5 | Complexes | 140 | 35 | 55 | 230 |
|  | Positive samples | 12,866<br>(9.09%) | 3,138<br>(0.2%) | 4,871<br>(0.1%) | 20,875<br>(0.3%) |
|  | Negative samples | 128,660<br>(90.9%) | 1,874,322<br>(99.8%) | 4,953,446<br>(99.9%) | 6,956,428<br>(99.7%) |
| Dockground | Complexes | 236 | 60 | 100 | 396 |
|  | Positive samples | 14,007<br>(9.09%) | 3,940<br>(0.05%) | 5,905<br>(0.04%) | 23,852<br>(0.11%) |
|  | Negative samples | 140,070<br>(90.9%) | 7,199,540<br>(99.94%) | 12,673,885<br>(99.95%) | 20,013,495<br>(99.88%) |
| MaSIF | Complexes | 1890 | 470 | 787 | 3147 |
|  | Positive samples | 308,441<br>(9.09 %) | 77,903<br>(0.28%) | 94,135<br>(0.26%) | 480,479<br>(0.74%) |
|  | Negative samples | 3,084,229<br>(90.9%) | 27,074,411<br>(99.71%) | 34,833,241<br>(99.73%) | 64,991,881<br>(99.26%) |

